# Complex spikes encode reward expectation signals during visuomotor association learning

**DOI:** 10.1101/841858

**Authors:** Naveen Sendhilnathan, Anna Ipata, Michael E. Goldberg

## Abstract

Climbing fiber input to Purkinje cells has been thought to instruct learning related changes in simple spikes and cause behavioral changes through an error-based learning mechanism. Although, this framework explains simple motor learning, it cannot be extended to learning higher-order skills. Recently the cerebellum has been implicated in a variety of cognitive tasks and reward-based learning. Here we show that when a monkey learns a new visuomotor association, complex spikes predict the time of the beginning of the trial in a learning independent manner as well as encode a learning contingent reward expectation signal after the stimulus onset and reward delivery. These complex spike signals are unrelated to and were unlikely to instruct the reward based signal found in the simple spikes. Our results provide a more general role of complex spikes in learning and higher-order processing while gathering evidence for their participation in reward based learning.

## Introduction

The cerebellum has been classically considered to be one of the centers for supervised learning in the brain, where the predicted results of movement are compared with the animal’s actual performance in relation to their sensory experience, in order to correct the errors in the action that led to the mismatch. The cerebellar cortex has been posited to achieve this via its two distinct types of inputs to its principle output cells, the Purkinje cells (P-cells). First, an efference copy of the ongoing plan generated by other areas in the brain, communicated to the P-cells through mossy fibers, and read out as high frequency simple spikes (SS); and second, a putative instruction signal, motioning unexpected events, communicated through the projections from the inferior olive, the climbing fibers, which evoke low frequency complex spikes (CS). The precisely timed relationship between the coincidence of complex spikes and simple spikes has been to shown to cause a long-term depression at the granule cell->P-cell synapse thereby supervising the information being learned at the level of P-cells^1^.

The CS have several characteristics: First, they fire with extremely low firing rates (baseline: 0.5-1 Hz, to ~10 Hz), second, they encode sparsely by only firing in about 20-30% of the trials^2,3^, third: some are predominantly evoked by mismatches in sensory predictions^4^, fourth, they can encode information in their rate of firing^2^, timing of firing^3^ or the duration of spikes^5^, fifth, presence of a single complex on a trial could potentially drive learning and might cause about 1 −10 sp/s change in the magnitude of simple spike firing rate in the next trial^3,5^. However, due to the combined effect of low frequency, sparseness and low amount changes that it causes to simple spike, significant changes in behavior, as a consequence of learning, only happens over several tens of trials. This flow of information and circuitry explains many simple motor learning behaviors from classical conditioning to motor adaption through gain changes: Connections that led to erroneous and unwanted behavior could be carefully pruned by the instructions provided by the complex spikes.

Recently, the cerebellum has been implicated in a variety of tasks that do not involve changes in the kinematics of movement but nevertheless have differences between the predicted consequences and the animal’s experience^6–10^. Cerebellum dependent learning of these behaviors cannot be readily explained by the present classical models. This is because, here, for example, rather than pruning connections that led to erroneous behavior, connections that would lead to preferred behavior need strengthening. Thus far, the role of CS in such reward processing has only been studied in classical conditioning, where they encode expectation of reward through a temporal difference prediction error framework^6–10^. The role of complex spikes in learning complex association tasks is still unknown.

Here, we studied the activity of the complex spikes recorded as a part of a previous data set ^11^ when monkeys learned to make associations between arbitrary visual cues and movements. We show that the CS encode a learning contingent reward expectation signal but did not instruct changes in simple spike dynamics. Classifying P-cell CS responses based on the SS characteristics revealed distinct mechanisms for reward-based learning in the mid-lateral cerebellum. Furthermore, in a paradigm with predictable temporal characteristics, CS exhibited a signal which anticipated the beginning of the next trial before the appearance of the cue and was unrelated to cell-type or the progression of learning. Our results provide a more general role of CS in learning and higher-order processing while gathering evidence for their participation in reward based learning. This while strengthening the current view that cerebellum is involved in reward-related, non-motor learning, adds critical constraints in the way cerebellum could achieve this function.

## Results

We trained two monkeys to perform a two-alternative forced-choice discrimination task ^11^, where, in each session, the monkeys learned to associate one of two novel visual symbols with a left-hand movement and the other visual symbol with a right-hand movement through trial and error. When the monkeys grabbed two bars with either hands, a small cue appeared on the screen immediately and the trial began. After a fixed duration (800 ms), one of the two symbols appeared on the screen and the monkeys released the hand associated with that symbol with a well learned-stereotypic hand movement to earn a liquid reward (delivered 1 ms after correct movement onset) as soon as possible (Fig 1a-b). The kinematics or the dynamics of hand movement were task irrelevant and only the release of the bars associated with the symbols merited reward. Monkeys usually performed a few trials (~30) of an overtrained association at the beginning of each session. Then, the monkeys were presented with novel symbols. They typically learned new associations (achieve criterion for learned) in ~50-70 trials on an average through an adaptive learning mechanism using a win-stay-lose-switch strategy (Fig 1c), similar to previous reports ^12^. Their reaction time generally was high during early learning and decreased significantly through learning (Fig 1c). The monkeys were free to move their eyes and thus occasionally made task irrelevant eye movements. The movement kinematics did not change at the symbol switch or through the progression of learning^13^.

**Figure 1:**
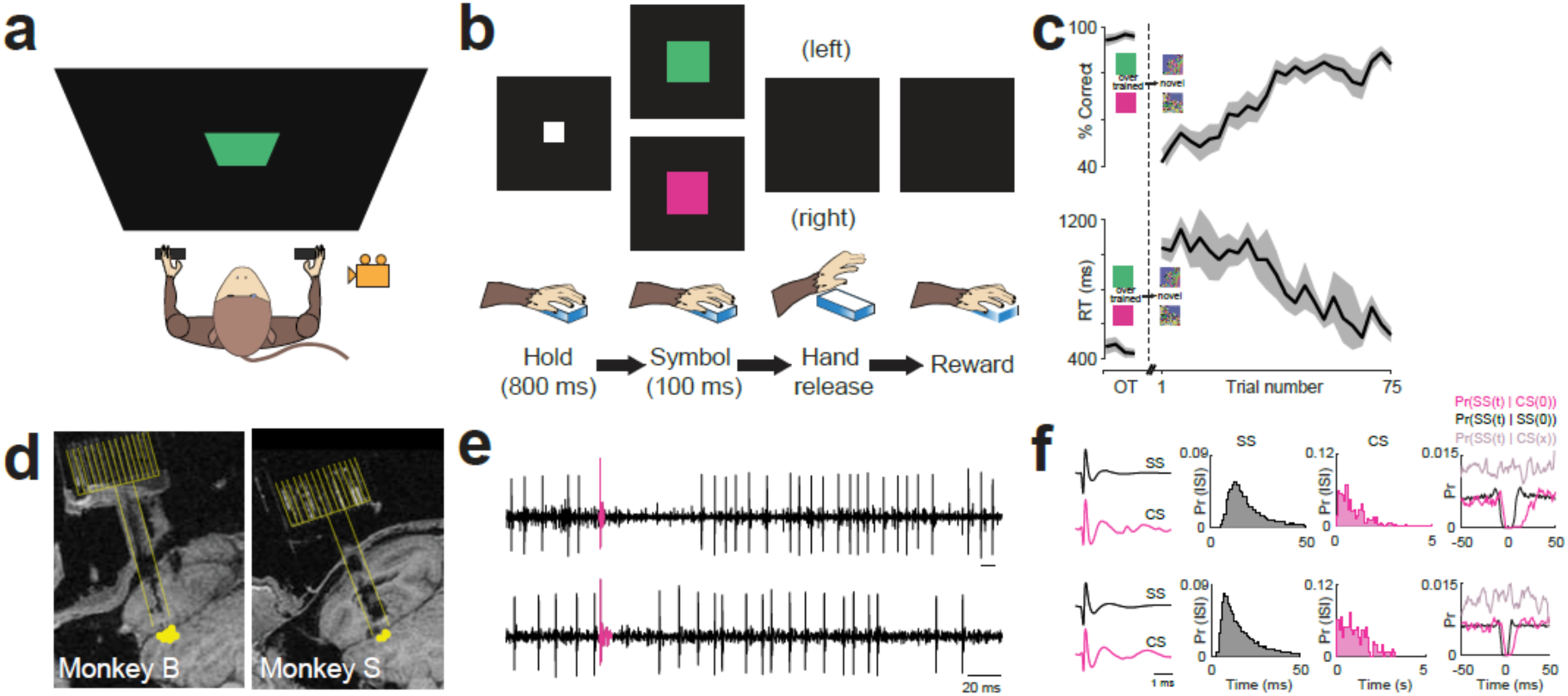
Two-alternative forced choice discrimination task, behavior and cerebellar electrophysiology. a. Schematic of the recording setup. b. The task started with the appearance of a white fixation spot when the monkey held on two manipulanda with either hands. Then one of the two novel visual symbols appeared briefly. The monkey then had to make a hand release associated with the presented symbol. Correct associations earn a drop of liquid reward immediately after the bar release onset. c. Mean learning curve (top) and mean reaction time (bottom) across all sessions. d. Coronal MRI of two monkeys (B and S) showing the single unit recording locations. e. Two representative raw neural signals from monkey B (top) and monkey S (bottom). f. Top panel: left: mean simple spike (SS) and complex spike (CS) waveforms from the cell shown in Fig 1e **top panel.** Middle: distribution of SS and CS interspike intervals (ISI) for the same cell. Right: Conditional probabilities of spike timings. Bottom panel: same as top panel but for cell shown in Fig 1e **bottom panel**.

We recorded near crus I and II of the mid-lateral cerebellum (Fig 1d). We identified P-cells by the presence of complex spikes online (Fig 1e), and offline by the i) spike waveforms (Fig 1f), ii) the SS and CS interspike interval distribution (Fig 1f) and iii) a pause in SS after a CS (Fig 1f, Fig S1) ^14^. We analyzed the SS for the cells that passed the above criteria (N=128 cells) in a previous study^13^. However, in this study, we only analyzed the CS from those cells that had reliably detected CS that were stable throughout the entire recording (N=25 cells).

While studying neural responses in the cerebellum, it is important to distinguish the possible interaction among three major components: SS activity, CS activity and behavior. This is because these three components are linked by fundamentally different mechanisms that operate independently on two levels: anatomically and temporally. Therefore, a comprehensive investigation would entail characterizing the effect of learning related changes in the SS modulation (which we discussed in detail elsewhere ^13^), learning related changes in CS responses and the effect of CS on the SS activity. Below, we discuss the latter two.

### Complex spikes responses during an overtrained visuomotor association task

First, we describe the CS responses when the monkeys were performing the overtrained task. We defined a trial to start from the reward information latency of SS (RIL, ~300 ms after reward onset, see ^13^) of one decision through the RIL of the consecutive decision. Based on the conventional role of CS in signaling unexpected events, we hypothesized that specific sensory events such as the symbol onset and or the reward delivery in the task might evoke a CS response in the P-cells. However, different P-cells showed modulations in CS firing rates in different epochs throughout the trial. Furthermore, many neurons showed changes in firing rates in more than one epoch (Fig 2a). Therefore, defining a ‘baseline epoch’ for performing any statistical analyses aimed at comparing firing rates was difficult. Consequently, to identify the epochs where the CS firing rate for a given neuron was statistically significant, we could not perform a simple statistical test comparing the baseline firing rate with the firing rate in the epoch of interest.

**Figure 2:**
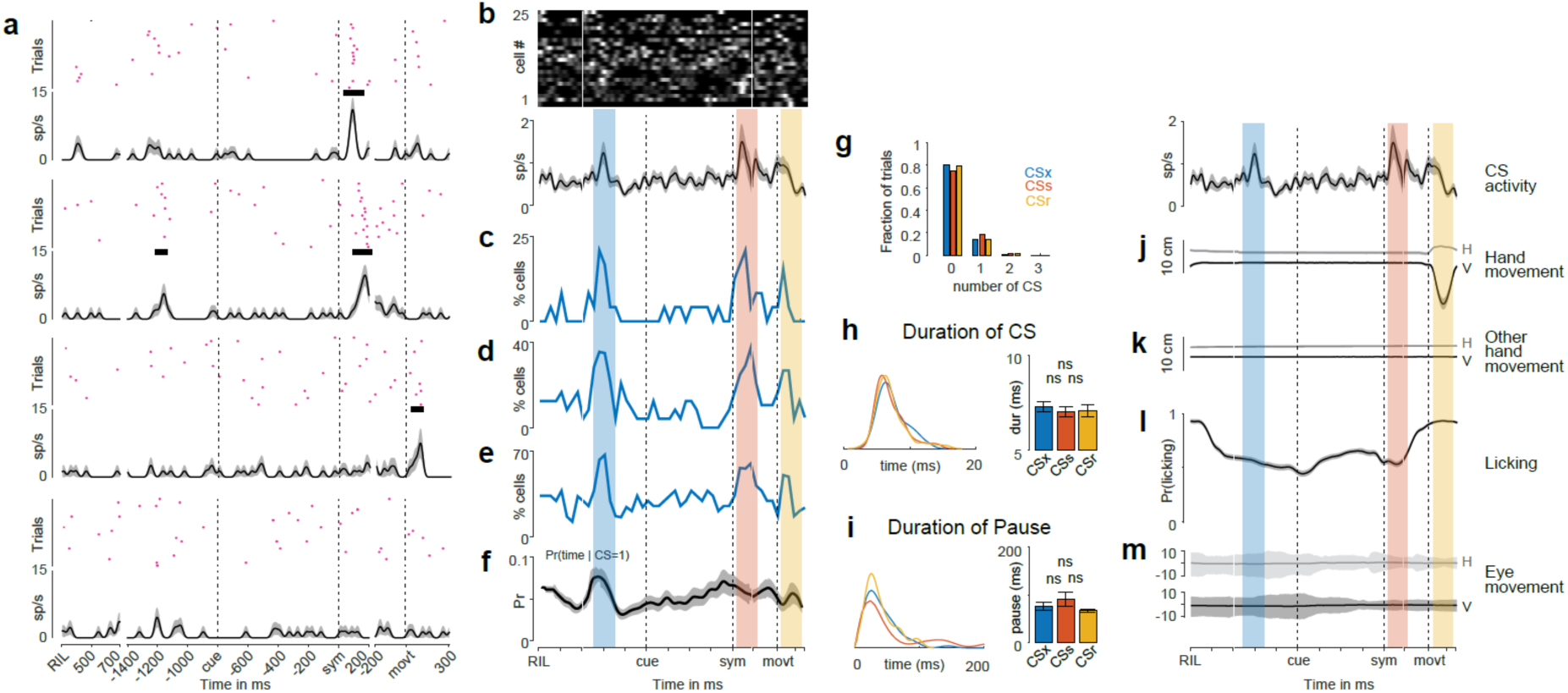
Complex spikes responses during a overtrained visuomotor association task. **a.** CS responses from four representative P-cells during the entire trial period (from one RIL to the next). Each of the top panels represent rasters and bottom panels represent spike density functions (sigma = 10 ms). The horizontal thick line on top of each spike density function represents the epoch where the activity was significant (estimated from the method shown in Fig S2a). **b.** Top: Heat map of CS responses from 25 P-cells (each horizontal line is a cell) during the entire trial period. Bottom: spike density function of population CS responses. **c.** Percent P-cells that showed significant change in neural activity during the trial period estimated from the method shown in Fig S2a. Blue, red and yellow shaded regions represent CSx, CSs and CSr epochs. These were the epochs in which the CS activities were analyzed for expectation, symbol and reward related activities. **d.** Same as above but estimated from the method shown in Fig S2b. **e.** Same as above but estimated from the method shown in Fig S2c. **f.** Conditional probability of CS timing when the CS was present. **g.** Fraction of trials with 0,1,2 or 3 CS in the CSx (blue), CSs (red) and CSr (yellow) epochs. **h.** Duration of CS waveforms in the CSx (blue) and CSs (red) epochs were not significantly different (P = 0.2460; ranksum test). Same was the case between CSr (yellow) and CSs (red) epochs (P = 0.2993; ranksum test) and between CSr (yellow) and CSx (blue) epochs (P = 0.8217; ranksum test). **i.** Duration of CS-SS pause in the CSx (blue) and CSs (red) epochs were not significantly different (P = 0.9719; ranksum test). Same was the case between CSr (yellow) and CSs (red) epochs (P = 0.0678; ranksum test) and between CSr (yellow) and CSx (blue) epochs (P = 0.1095; ranksum test). **j.** Mean horizontal (gray) and vertical (black) positions of the responding hand. **k.** Mean horizontal (gray) and vertical (black) positions of the non-responding hand. **l.** Mean licking behavior **m.** Mean horizontal (gray) and vertical (black) eye positions. All shadings represent s.e.m unless stated otehrwose.

We developed a method to compensate for this problem (see methods; Fig S2a). We found that 94% of cells showed at least one epoch of significant activity. At the population level (Fig 2b), we found that P-cells tended to change their firing rates in three distinct epochs: 500-300 ms before the cue onset (called CSx epoch; ~24% of cells), 50-250 ms after the symbol onset (called CSs epoch; ~23% of cells) and 50-250 ms after the reward onset (called CSr epoch; ~20% of cells) (Fig 2c). We confirmed this result using two additional analyses that differed in the amount of stringency in classifying neurons as showing significant activity in a given epoch (see methods, Fig 2d-e; Fig S2b-c).

The timing of CS in the CSx epoch, calculated as the conditional probability of time (in this epoch) given there was a complex spike, i.e., Pr(time | CS = 1), was significantly high (P = 0.0002; ranksum test) indicating the high temporal precision in CS timing (Fig 2f), peaking at ~400 ms before the cue. Therefore, we reasoned that given the regularity of the time of appearance of the cue after the hand movement of the previous trial, the CS activity in this epoch was a predictive timing signal, predicting the appearance of the cue rather than a reactive signal triggered by any task-related event. The CS fired in about 18% of trials in this epoch consistent with prior reports of the probability of CS responses ^2,3^ (Fig 2g).

There were also CS responses in the CSs and CSr epochs linked to the stimulus appearance and reward respectively. The CS responded significantly in about 21% of trials in the CSs epoch and in 19% of trials in the CSr epoch (Fig 2g). This suggest that the CS did not provide a temporal-difference prediction error, which would have predicted little or no response in the CSr epoch^6^. Furthermore, we did not see any modulation in CS duration (Fig 2h; P= 0.4017, ANOVA) or the duration of pause in SS firing elicited by a CS (Fig 2i; P = 0.1670; ANOVA) among these three epochs. The CS responses in any of the three epochs could not be explained by any obvious changes in motor kinematics, such as movement of the responding hand (Fig. 2j), the non-responding hand (Fig 2k), licking^8–10^ (Fig 2l) or eye movements (Fig 2m). Henceforth, we only analyze the CS activity in these three epochs.

The CS responses in the CSs and CSr were not selective for hand or symbol. We first calculated the contrast function (A-B)/(A+B) in the symbol epoch (50-250 ms after symbol onset) for preferences between the two symbols and in the movement epoch (50 ms before to 250 ms after the movement onset) for preferences between the hand movements and the symbols. To verify if this tuning was meaningful and not just due to extreme differences in sampling number and noise, we generated a null distribution of spike times through a gamma distribution that was matched with the parameters of the experimental data (we obtained the shape parameter the ISI distribution fit and took the scale parameter as the inverse of firing rate) and calculated a similar tuning function on this null distribution. We found that the CS responses during the symbol (Fig 3a) or the movement epochs (Fig 3b) were not statistically different from a null distribution (Symbol selectivity: P = 0.5153; t-test; Movement selectivity: P = 0.4811; t-test).

**Figure 3:**
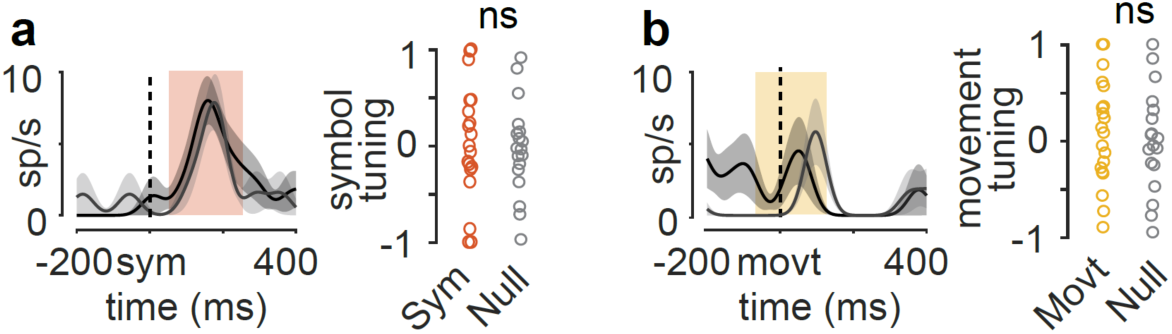
CS responses were not tuned to sensorimotor parameters. a. Left: CS responses from a representative P-cell for symbol 1 (gray) and symbol 2 (black). Shaded region shows the epoch where the CS activity was analyzed. Right: symbol tuning index for the P-cells (red) was not different from that of a null population (gray) (P = 0.5153, t-test). See methods for more details. b. Left: CS responses from a different representative P-cell for left (gray) and right (black) hand release. Shaded region shows the epoch where the CS activity was analyzed. Right: movement tuning index for the P-cells (yellow) was not different from that of a null population (gray) (P = 0.4811, t-test). See methods for more details.

### Cell type specific CS signals provided a learning-dependent reward expectation signal

Next, we studied the learning related changes in CS response in both symbol and the reward epochs (CSs and CSr). We previously showed that the mid-lateral cerebellar P-cells SS encode a reward-based error signal when monkeys learn a new visuomotor association, ^13^ reporting the outcome of the most recent decision in short epochs called ‘delta epochs’^13^. The error signal in the delta epoch neither described an error in the parameters of the motor effector used to report the choice (hand) nor any other task-irrelevant movements (such as eye movements, licking or swallowing), nor errors in sensory parameters such as the sound of the solenoid click, the symbols used to for association etc. During learning, roughly half of the P-cells were selective for correct outcome (cP-cells) and the remaining were selective for wrong outcome (wP-cells). As the monkey learned the associations, the difference in activity after successful and unsuccessful trials decreased through a reinforcement learning framework.

Therefore, we looked at the learning related changes in CS responses in cP-cells and wP-cells separately through a reinforcement learning framework. To do this, we analyzed the CS activity in during three phases of learning: First 20 trials (early learning trials, L1), middle 20 trials (mid-learning trials, L2), 20 trials after reaching the criteria for learned (learned trials, L3) (Fig 4a, Fig S3). We hypothesized that if the CS encoded reward-related activity, then they should show learning related changes in the symbol and the reward epoch. We found significant learning related changes in CS responses for both types of P-cells, although in very different ways as described below (Fig 4b, 4e).

**Figure 4:**
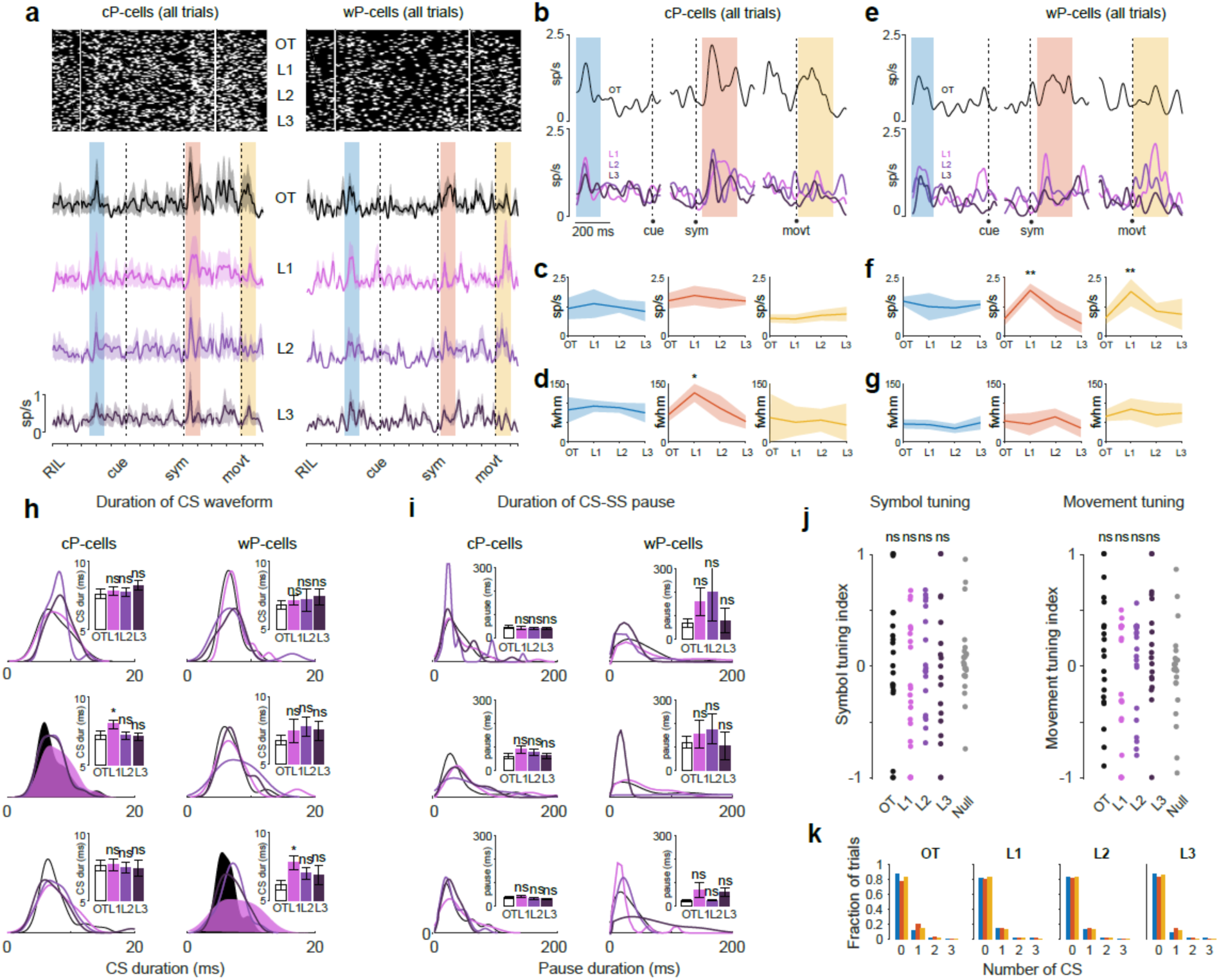
Cell type specific CS signals provided learning dependent reward expectation signal. a. Top left: Raster for all the cP-cells for the entire trial duration for OT, L1 L2 and L3 conditions. Bottom left: spike density functions for OT, L1 L2 and L3 conditions. Blue, red and yellow shaded regions represent CSx, CSs and CSr epochs respectively Right: Same as left but for all wP-cells. b. Top panel: spike density functions of cP-cells in the OT condition for CSx (left), CSs (middle) and CSr (right) epochs. Bottom panel: spike density functions of cP-cells in L1, L2 and L3 conditions for the same three epochs. c. Peak firing rate of cP-cells in OT, L1 L2 and L3 conditions for CSx (left), CSs (middle) and CSr (right). d. Temporal dispersion of CS activity for cP-cells (estimated as the full width at half maximum firing rate) in OT, L1 L2 and L3 conditions for CSx (left), CSs (middle) and CSr (right). e. Same as **b** but for wP-cells. f. Same as **c** but for wP-cells. g. Same as **d** but for wP-cells. h. Left: duration of CS waveforms for cP-cells in OT, L1, L2 and L3 conditions for CSx (top), CSs (middle) and CSr (bottom) epochs. Right: same as left but for wP-cells. i. Left: duration of CS-SS pause for wP-cells in OT, L1, L2 and L3 conditions for CSx (top), CSs (middle) and CSr (bottom) epochs. Right: same as left but for wP-cells. j. Left: Symbol tuning index for OT, L1, L2 and L3 conditions and null population. Right: same as left but for movement tuning index. k. Fraction of trials with 0,1,2 or 3 complex spikes in OT, L1, L2 and L3 conditions. * means P<0.05; ** means P<0.01.

For cP-cells, after the symbol switch, the peak firing rate of CSs did not change between the OT and learning trials (P=0.6344; ranksum test, Fig 4c). However, the CSs activity trended toward being temporally more dispersed (estimated as the full width at half maximum firing rate, fwhm) during early learning compared to OT condition (P=0.0234; ranksum test Fig 4d). We suggest that this is because the monkeys performed at chance level during early learning (Fig 1c) and thus the new symbols were no longer reliable predictors of upcoming reward as they were in the OT condition. However, over the course of learning, as the monkeys learned the association between symbols and the movements, the CSs activity became temporally less dispersed (i.e. more temporally precise) as the symbols more accurately represented a future reward (P = 0.3421; ranksum ttest; Fig 4c). Therefore, the CSs of cP-cells encoded a neural correlate of reward expectation during visuomotor association learning (Fig4b-d).

For wP-cells, the peak firing rate of CSs, and not the temporal dispersion, reflected the learning state of the animal (Fig 4e-g): CSs increased their firing rate during early learning (P=0.0135; ranksum test) and systematically decreased their firing rate with learning (P=0.5294; ranksum test; Fig 4f). Additionally, unlike the cP-cells, the wP-cells also showed learning related changes in activity in the CSr epoch. During early learning, the CSr activity was higher and after a delay of ~100 ms^2^. Since the Pr(reward) ~ 0.50 during early learning, the CSr activity only increased after experiencing the reward delivery (or its absence) which is ~100 ms. This is because the experience of reward (or its absence) acted as an unexpected event whose information was available only after its experience^6^. In the second third of learning, as the Pr(reward) increased, the CSr response decreased in magnitude and shifted earlier, closer to the movement onset. In the last third of learning, when the Pr(reward) ~ 1, not only the CSr response was almost at the spontaneous level, it also occurred immediately after the movement onset (Fig 4e). In summary, although the familiarity of symbols served as an indicator of the expected reward, the monkeys had to make the correct choice of hand release given a symbol to earn the reward. Therefore, as the Pr(reward) increased during learning, the CSr activity was paired strongly with the movement preparation. After learning, as the movement was being prepared, the CSr were also fired simultaneously.

Furthermore, the duration of CS was different between the learning conditions, also in a cell type dependent way. As discussed above, the cP-cells showed changes in CS firing rate only in the symbol epoch (Fig 4b-d). Consistent with this, we found that, only in the symbol epoch, the CS waveform was longer during the early learning period compared to OT condition (P = 0.0379 ranksum test; Fig 4h) and progressively became shorter through learning, finally resembling the waveform in the OT condition in duration (P = 0.7822 ranksum test; Fig 4h). The CS duration for cP-cells did not change in the reward epoch during learning (P = 0.6496, ranksum test; Fig 4h). However, there was no learning related changes in the CS-SS pause in either for CSs (P = 0.2220, ranksum test; Fig 4i) or CSr (P = 0.5358, ranksum test; Fig 4i). Additionally, the CS activity was not selective for symbols (P = 0.4343; ANOVA; Fig 4j) or the choice of hand (P = 0.5404; ANOVA; Fig 4j) during learning.

For wP-cells, the duration of CS was longer during the early learning period compared to OT condition (P = 0.0242 ttest; Fig 4h) only in the reward epoch. The duration progressively trended towards being shorter through learning, (P = 0.3352, t-test; Fig 4h). The CS duration for wP-cells did not change in the symbol epoch during learning (P = 0.2650, t-test; Fig 4h). Again, there were no learning related changes in the CS-SS pause for CSs (P = 0.4963, ranksum test; Fig 4i) or CSr (P = 0.7515, ranksum test; Fig 4i). Additionally, the CS activity for wP-cells also continued to lack any tuning for symbols (P = 0.3757; ANOVA; Fig 4j) and movement (P = 0.1752; ANOVA; Fig 4j) through learning. Taken together, we found that the CS responses varied significantly for cP-cells and wP-cells although they both encoded learning dependent reward expectation signal.

### P-cells encoded learning and cue independent expectation signal

In addition to the reward expectation signals in the CSs and CSr epochs, P-cells encoded a very different expectation signal in the CSx epoch that did not depend the cell-type. Just as in the OT condition, the CSx activity signaled a learning independent and cue independent preparatory signal for the beginning of the trial (Fig 2f, Fig 4a-g). This suggests the presence of an internally generated signal for the expectation of the task that is independent of any form of cue but is only dependent on ‘time’ as the only cue.

Although the CSx activity signaled an expected beginning of a trial, it was learning independent. That is for either type of P-cells, neither the peak activity (cP-cells: P =0.2458, ranksum test; Fig 4c; w-cells: P =0.3595, ranksum test; Fig 4f), the temporal dispersion in activity (cP-cells: P =0.8902, ranksum test; Fig 4d; w-cells: P =0.7349, ranksum test; Fig 4g), CS waveform duration (cP-cells: P =0.6641, t-test; Fig 4h; w-cells: P =0.2650, t-test; Fig 4h) or the CS-SS pause (cP-cells: P =0.7431, t-test; Fig 4i; w-cells: P =0.4597, ranksum test; Fig 4i) changed with learning. Both types of P-cells showed this signal precisely at the same time (Fig 4b, e). Therefore, we propose that this signal purely represents a task relevant, learning independent expectation signal.

### Complex spike activity was unrelated to changes in simple spike activity

We previously showed that although the delta epochs occurred at different times across the population, from immediately after the RIL of the prior trial to just before the RIL of the current trial they were consistent across trials for a given P-cell. The information about the most recent decision collectively spanned the entire trial period. To investigate the relationship between complex spikes and simple spikes, we asked if there were any temporal relationship between CS activity and the delta epoch of SS. First, if the changes in the simple spike neural activity in delta epoch were caused by complex spikes, we should either expect a tiling effect of complex spikes similar to what we found for the simple spikes^13^ or a modulation of complex spike activity after the feedback containing information about the trial outcome which could then induce delta epochs with varying latencies in simple spike through an unknown mechanism. However, we have four lines of evidence against this hypothesis: First, both types of P-cells lacked information about the prior trial outcome in any of the three epochs: CSr, CSs or CSx (cP-cell: Fig 5a-b; CSx: P = 0.7999, paired t-test; CSs: P = 0.5021, paired t-test; CSr: P = 0.6338, paired t-test and wP-cell: Fig 5c-d; CSx: P = 0.1524, paired t-test; CSs: P = 0.8815, paired t-test; CSr: P = 0.1064, paired t-test). Second, the time of modulation of CS activity during learning for a given P-cell was not correlated with any fixed time relative to delta epoch. The time of modulation of CS activity during learning was not clustered around the start of the delta epoch (cP-cells: P = 0.33, Fig 6c; wP-cells: P = 0.30, Fig 6d Rayleigh test for non-uniformity of circular data), middle of the delta epoch (cP-cells: P = 0.50, Fig 6c; wP-cells: P = 0.09, Fig 6d Rayleigh test for non-uniformity of circular data), or the end of the delta epoch (cP-cells: P = 0.73, Fig 6c; wP-cells: P = 0.39, Fig 6d Rayleigh test for non-uniformity of circular data). The time of CS modulation in the OT condition did not correlate with the time of delta epoch in the learning condition either suggesting that the afferent CS activity did not instruct the formation of the delta epoch in the simple spikes (start of the delta epoch: cP-cells: P = 0.82, Fig 6c; wP-cells: P = 0.93, Fig 6d; middle of delta epoch: cP-cells: P = 0.89, Fig 6c; wP-cells: P = 0.96, Fig 6d; end of delta epoch cP-cells: P = 0.91, Fig 6c; wP-cells: P = 0.94, Fig 6d Rayleigh test for non-uniformity of circular data). Third, unlike the simple spikes, the CS activity at the start, middle or the end of delta epoch during learning did not carry information about the prior trial outcome ^2^ (Fig 6e start of the delta epoch: cP-cells: P = 0.4360 paired ttest, wP-cells: P = 0.5189, paired ttest; middle of delta epoch: cP-cells: P = 0.4106, paired ttest wP-cells: P = 0.8463, paired ttest; end of delta epoch cP-cells: P = 0.3708, ranksum test, wP-cells: P = 0.9899 ranksum test). Finally, P-cells with the same CS responses had very different delta epochs (Fig S3a); P-cells with similar delta epochs had very different CS responses (Fig S3a) and furthermore, some P-cells with delta epochs did not show any modulation in CS, suggesting a dissociation between the two (Fig S3a). All these provide strong converging evidence that complex spikes were unlikely to instruct a change in simple spike activity through the classical error-based learning framework.

**Figure 5:**
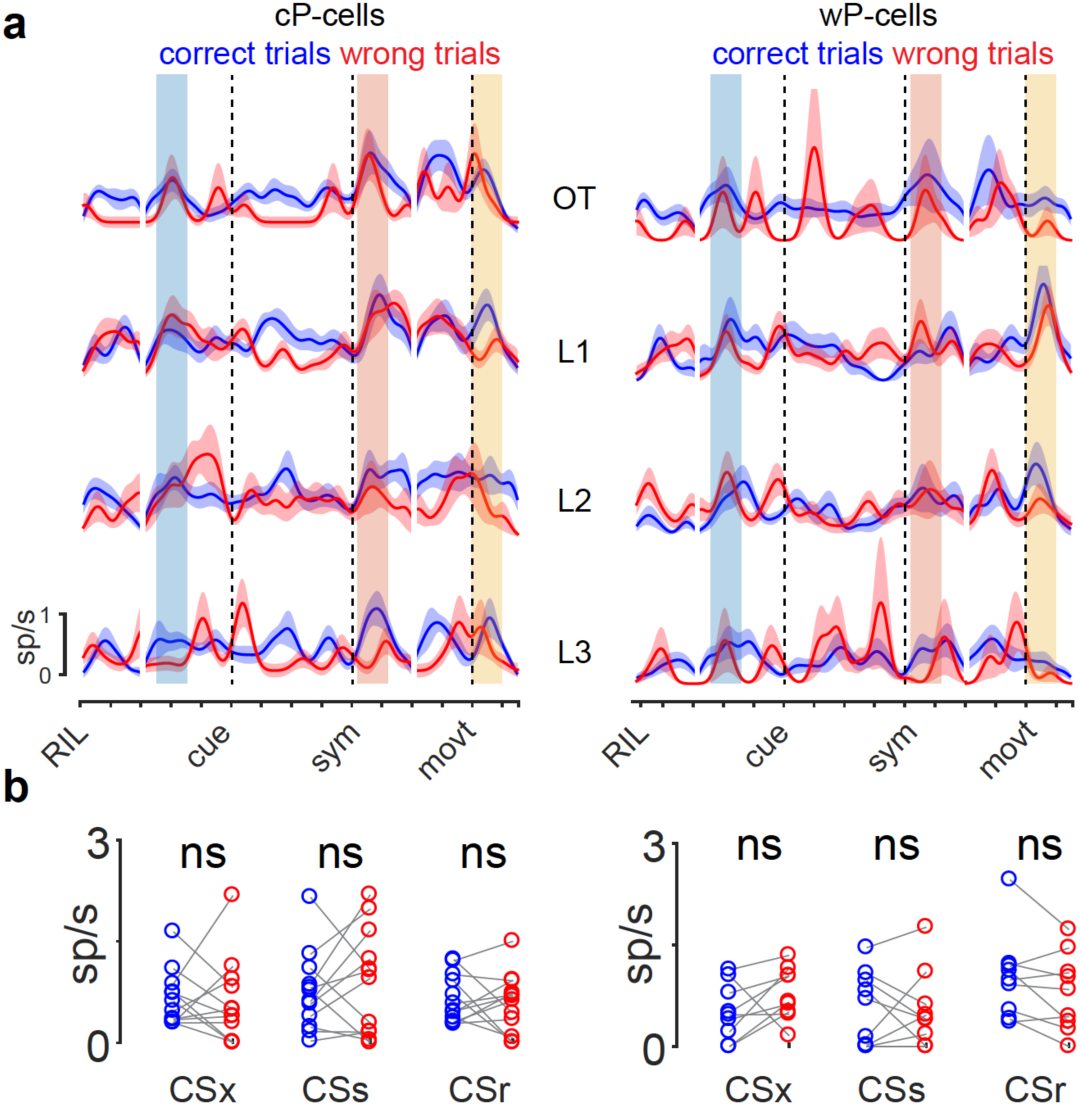
Complex spike did not carry information regarding trial outcome. a. Same as Fig 5a but separated into correct (blue) and wrong (red) trials. b. Quantitation of activity responses from a for cP-cells (left; CSx: P = 0.7999, paired t-test; CSs: P = 0.5021, paired t-test; CSr: P = 0.6338, paired t-test) and wP-cells (right; CSx: P = 0.1524, paired t-test; CSs: P = 0.8815, paired t-test; CSr: P = 0.1064, paired t-test).

**Figure 6:**
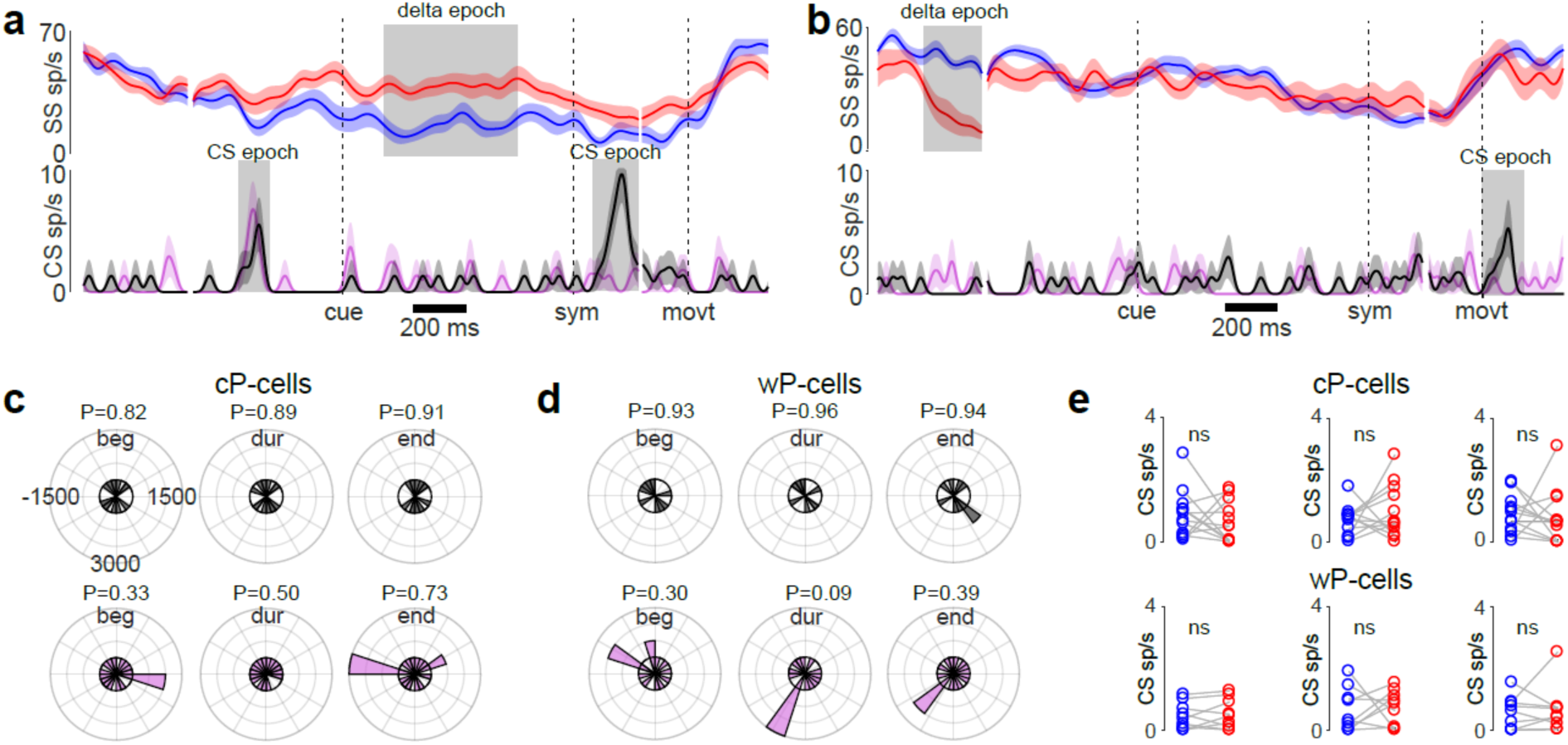
Complex spike activity was unrelated to changes in simple spike activity. a. Top: Representative wP-cell simple spike activity during learning for correct (blue) and wrong (red) trials. Shaded region is the delta epoch. Bottom: CS activity from the same P-cell during OT condition (black) and learning (pink). Shaded region is the epoch in which CS was modulated significantly. b. Same as a, but for a cP-cell. c. Circular histogram (of the entire trial period) of CS modulation times in OT (top) and learning (bottom) relative to the cell’s time of beginning (left), center (middle) or end (right) of the delta epoch during learning for all cP-cells. Values on the plot represent time in ms from the trigger (beginning, center or end of delta epoch for the appropriate plot). d. Same as above but for wP-cells. e. Peak CS activity for correct (blue) and wrong (red) trials during learning at the start (left), during (middle) or the end (right) of delta epoch for cP-cells (top) and wP-cells (bottom). Start of the delta epoch: cP-cells: P = 0.4360 paired ttest, wP-cells: P = 0.5189, paired ttest; middle of delta epoch: cP-cells: P = 0.4106, paired ttest wP-cells: P = 0.8463, paired ttest; end of delta epoch cP-cells: P = 0.3708, ranksum test, wP-cells: P = 0.9899 ranksum test.

## Discussion

The role of complex spikes in the functioning of the cerebellum is not well understood^15^. Different studies, using different behavioral paradigms in different species have come up with a number of different and seemingly contradictory suggestions for the role of complex spikes^15^. CS activity across different cerebellar regions of the same animal doing the same task could also be vastly different^8^.

The classic Marr-Albus model for error-based motor learning posits that the coincidence of complex spike firing and the parallel fiber input leads to synaptic change at the granule cell-to-P-cell synapses, facilitating trial-by-trial correction of motor errors. In this case, CS activity reports an error, and not necessarily a description of the action itself. CS can encode error magnitude in its probability of firing^2^, its timing^3^ or its waveform duration^5^ in different or even in the same cerebellar area. In contrast, in the oculomotor system, for smooth pursuit^16^ and the otololithic vesitibulo-ocular reflex^17^, CS activity describes the movement itself, with CS inhibition of simple spikes providing a reciprocal modulation in the simple spike activity. In well-learned reaching tasks in the monkey, the CS of different P-cells report kinematic errors in position, velocity, and acceleration^18^

There is recent overwhelming evidence that the cerebellum is implicated in tasks beyond simple motor learning^8,10,13,19,20^. McCormick et al. ^21^ originally showed that the cerebellum is necessary for the Pavolvian association task of eye-blink conditioning, and since then, several others have shown that in eye-blink conditioning^6^ and in other forms of simple classical conditioning^8,9^ in the mouse, CS activity provides a temporal-difference prediction error. In a task where mice had to move a virtual wheel into a reward zone, the CS of P-cells in different zones of lobule simplex were excited or suppressed by reward, or the omission of reward^10^.

We previously showed that P-cell SS in the mid-lateral cerebellum transiently reported the result of the monkey’s most recent decision during visuomotor association learning in brief epochs, where half of the P-cells were excited and the rest were inhibited by a correct outcome and vice versa for a wrong outcome^13^. The SS activity changed with learning through a reinforcement learning framework. Here we analyzed the concurrent CS activity in the same task. We found striking patterns of CS activity, which could not be explained by any previous model of CS activity. We found CS firing above the background activity rate in three different epochs: ~400 ms before the cue that initiated the trial (CSx), in the period after the symbol appeared (CSs), and in the period after the reward (CSr) (Fig 2). Surprisingly, the CSx response occurred in both the overtrained and learning contexts, for all cells. Because this response occurred before the cue, it must have arisen from an intrinsic timing mechanism, and may have served as an alerting mechanism. This activity could not be explained by any changes in motor kinematics (hand, licking or eye movements) either. No other study, according to our knowledge, has shown such a preparatory signal. In contrast, the CSs responses changed only during the learning period. For cP-cells, the response did not decrement, but the response duration shortened as the monkey learned the task. For wP cells the response decremented, but the duration did not change. Both these learning related changes could signal a confidence in the probability of reward, from chance to certainty (Fig 4). The cP cells did not respond during the CSr epoch, unlike the wP cells, which responded during early learning with a large response whose rate decremented as the monkey learned. The CSr epoch response was not a simple reward response, since it occurred after failed as well as successful decisions (Fig 5). The changes in CS activity we report here were unlikely to have been due to any changes in sensory or motor events. This is because, we previously showed that the hand movements between correct and wrong trials and between the OT and learning remain the same. Other task irrelevant motor behavior such as licking or eye movements were also uncorrelated with the CS activity (Fig 2).

There are several reasons why the CS activity we have measured cannot fit a classical temporal-difference prediction error (TD) model. First, such a model requires that reward information be transferred from the actual reward event to the earliest reward predicting event. However, the activity in the earliest response epoch, the CSx, was entirely unrelated to actual trial events beyond the timing of the trial itself, and the greatest response in the symbol epoch, which should develop in a TD model as learning proceeds, occurred at the symbol switch, where in a TD model there should be no activity at all.

Despite the onset of a visual cue instructing a future reward, a correct choice of the hand was required for reward delivery. This action was therefore a necessary predictor of trial outcome. Consequently, we saw learning related changes in CS activity for both symbol and reward events, although in different ways for different cell types. We speculate that observed differences in CS activity between cell types could be due to the different sensorimotor input/output properties. Interestingly, similar signals have been shown in the dopamine system where the release of dopamine of contingent on the initiation of the correct action and not just the reward prediction^22^. A recent study in mice showed that the deep cerebellar nuclei (DCN) projects to the dopamine neurons in the VTA via a monosynaptic connection and is sufficient to drive social behavior^23^. Other studies have shown that in primates the cerebellum has a closed loop connection with the basal ganglia^24^. Although suggestive, our results in this study about the CS signals providing a reward expectation signal during learning combined with the results of our previous study^13^ about the SS signals providing a reinforcement learning contingent error signal fits well in the framework that the cerebellum might be working in tandem with the basal ganglia in effectively driving reward-related behavior.

Furthermore, we did not find any relationship between the CS activity and the SS activity^7,25^. One might have expected that a CS signal could have served as a teaching signal for the delta epoch of SS during learning if the classical error correcting framework were to apply. This was not at all the case (Fig 6). There are several reasons why complex spike signals are unlikely to play the role of a teaching signal in our experiment^20^. First, it is unlikely that the difference in the simple spike rate of more than 30 sp/s in the delta epochs between consecutive correct and wrong trials could be caused solely by synaptic depression elicited by complex spikes which has only been shown to cause a maximum of 8-10 sp/s changes in SS activity (with the longest CS waveforms) ^2,5^. In addition, if the complex spikes were causing the delta epochs, we should have seen a tight temporal relationship between the two, but we did not. From a mechanistic point of view, a complex spike signal has been shown to induce LTP which weakens the connections that led to unwanted behavior ^1,26,27^. This is more suited to error-based learning. However, in a reinforcement learning, rather than penalizing unwanted behavior, desirable behaviors are rewarded and those connections are strengthened ^28^. This again makes is unlikely for CS to instruct changes in SS firing. These observations suggest that the computations related to changes in activity in the delta epoch are performed within the cerebellum from a mechanism not involving complex spikes.

Taken together, our study is part of a rapidly growing literature body suggesting new and general roles for CS signals that are disparate from classical error-based supervised learning. We report two different CS signals when the monkey learns a visuomotor association. The first is a timing signal that could prepare the network for a new trial in anticipation of the cue. The second is not related to the presence or absence of reward, but could provide an estimate of the probability of success. How this probability estimate can facilitate learning remains to be seen.

## Methods

We performed all experiments on two adult male rhesus monkeys using methods and techniques that have been described in detail previously^11,13^. All experimental protocols were approved by the Animal Care and Use Committees at Columbia University and the New York State Psychiatric Institute, and complied with the guidelines established by the Public Health Service Guide for the Care and Use of Laboratory Animals. Briefly, we used the NEI REX-VEX-MEX system for behavioral control^29^. The monkey sat inside a dimly lit recording booth, with his head firmly fixed, in a Crist primate chair, 57 mm in front of a back-projection screen upon which visual images were projected by a Hitachi CP-X275 LCD projector controlled by a Dell PC running the NIH VEX graphic system.

### Two-alternative forced-choice discrimination task

The task began with the monkeys grasping two bar-manipulanda, one with each hand, after which a white square appeared for 800 ms. Then one of a pair of fractal symbols, that the monkey had never seen before, appeared briefly for 100 ms, at the center of gaze. One symbol signaled the monkey to release the left bar and the other to release the right bar. We rewarded the monkeys with a drop of liquid juice reward for releasing the hand associated with that symbol. Over 50 to 70 trials, the monkey gradually learned which symbol was associated with which hand.

### Single unit recording

We used two recording cylinders, on the left hemisphere of each monkey. We introduced glass-coated tungsten electrodes with an impedance of 0.8-1.2 MOhms (FHC) into the left mid-lateral cerebellum of monkeys every day that we recorded using a Hitachi microdrive. We passed the raw electrode signal through a FHC Neurocraft head stage, and amplifier, and filtered through a Krohn-Hite filter (bandpass: lowpass 300 Hz to highpass 10 kHz Butterworth), then through a Micro 1401 system, CED electronics. We used the NEI REX-VEX system coupled with Spike2 (CED electronics) for event and neural data acquisition. We verified all recordings off-line to ensure that we had isolated Purkinje cells and that the spike waveforms had not changed throughout the course of each experiment.

### Methods to detect epochs of significant changes in signal

#### Method1

We considered every 50 ms epoch as a ‘putative baseline epoch’ and performed pairwise t-tests comparing the CS activity in the putative baseline epoch with the CS activity in every other 50 ms epoch throughout the trial. This gave us a series of P-values for each epoch, testing the null hypothesis that the activity in the putative baseline epoch was statistically similar to the activity in every other epoch. Those epochs that were significantly different (P<0.05 after correcting for multiple comparisons) from at least 75% of the putative baseline epochs were considered to have a significant change in CS activity (Fig S2a).

#### Method2

Consider the CS activity from a 50 ms epoch of one trial as a ‘bin’. We performed a t-test between 50 bootstrapped bins from a given 50 ms epoch and 50 bootstrapped bins from all the epochs across the entire trial period. This gave as a P-value for that epoch, testing the null hypothesis that the activity in that epoch was statistically similar to the activity in any other epoch. We repeated this for every 50 ms epoch and found the epochs that had a P value < 0.05 (Fig S2b). ***Method3***: We considered −100 to 0 ms aligned on cue onset as the chosen baseline epoch and compared the activity in that epoch with every other epoch using a conventional approach. The epochs with P<0.05 were considered to have significantly different firing rate from the ‘baseline’ rate (Fig S2c).

#### Statistics

To check if two independent distributions were significantly different from each other, we first performed a two-sided goodness of fit Lilliefors test, to test for the normality, then used an appropriate t-test; or else a non-parametric Wilcoxon ranksum test. All values in this study, unless stated otherwise, are mean ± s.e.m.

## Acknowledgments

We thank Glen Duncan for highly creative and wonderful electronic assistance, John Caban and Matthew Hasday for superb machining, Dr. Girma Asfaw, Dr. Moshe Shalev for animal care and Lisa Kennelly, Whitney Thomas and Holly Cline for facilitating everything.

## Funding

This work was supported by the Keck, Zegar Family, and Dana Foundations and the National Eye Institute (R24 EY-015634, R21 EY-017938, R21 EY-020631, R01 EY-017039, and P30 EY-019007 to M. E. Goldberg, PI).

## Competing interests

Authors declare no competing interests.

## Data and materials availability

All data is available in the main text or the supplementary materials. Raw data and codes are available upon reasonable request.

## Disclosure

The authors declare no competing financial interests

---

**Figure S1:**
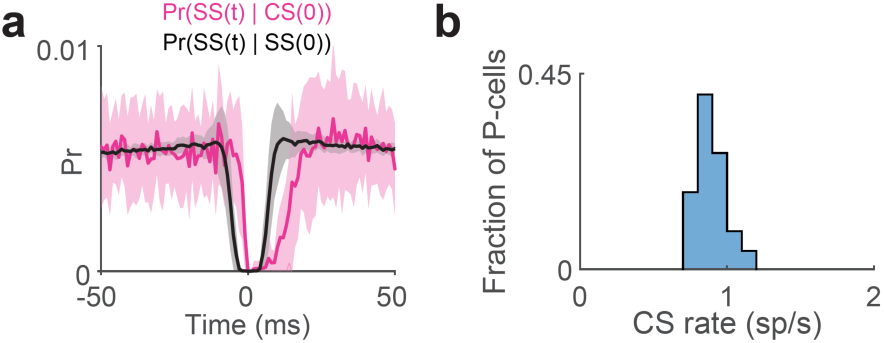
Complex spike properties. a. Conditional probability of time of simple spike at time t given there was a complex spike at time t=0 (pink) and the conditional probability of time of simple spike at time t given there was a simple spike at time t=0. b. Population CS rate histogram.

**Figure S2:**
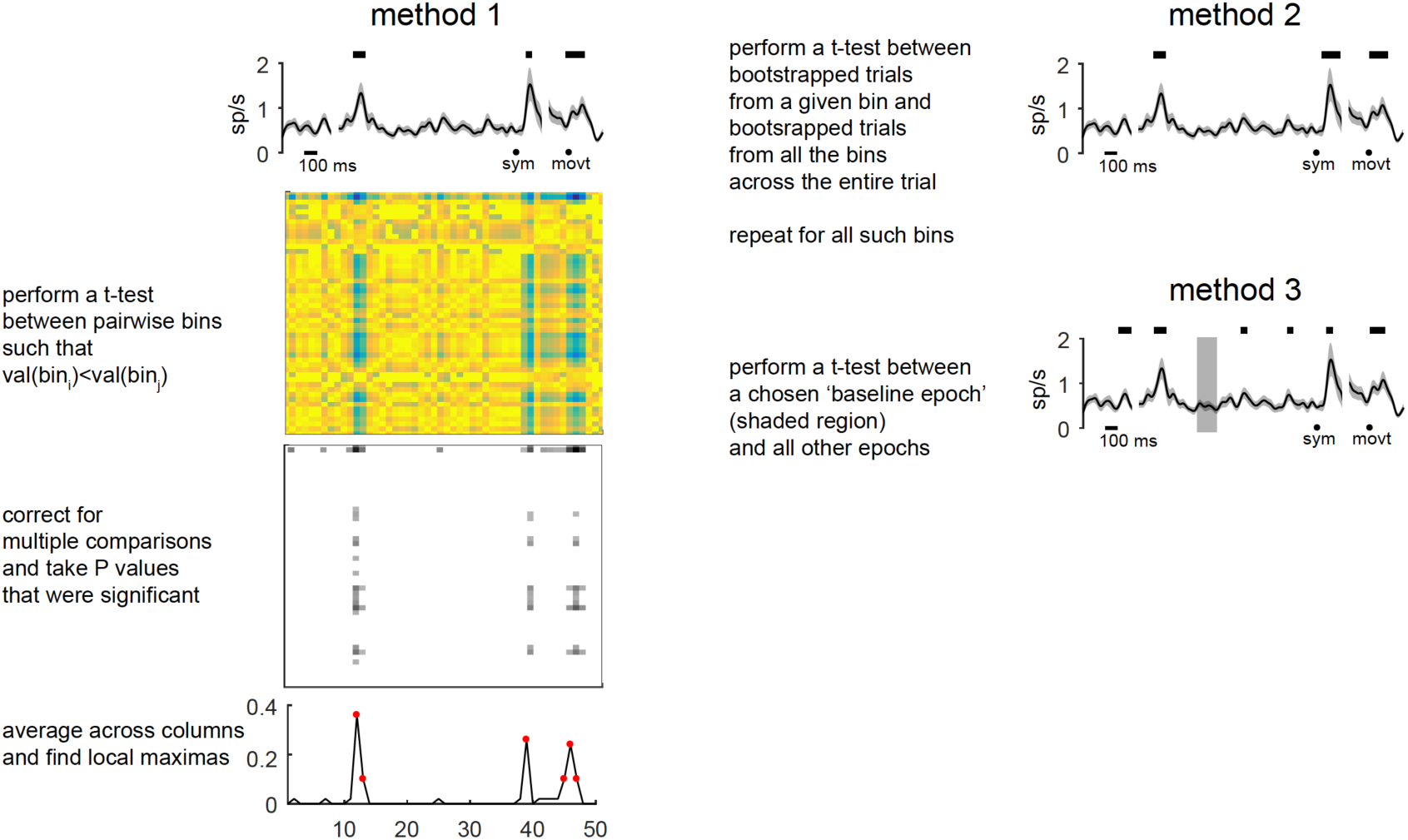
Methods to detect the epochs where the CS activity was significant. ***a. Method1***: We considered every 50 ms epoch as a ‘putative baseline epoch’ and performed pairwise t-tests comparing the CS activity in the putative baseline epoch with the CS activity in every other 50 ms epoch throughout the trial. This gave us a series of P-values for each epoch, testing the null hypothesis that the activity in the putative baseline epoch was statistically similar to the activity in every other epoch. Those epochs that were significantly different (P<0.05 after correcting for multiple comparisons) from at least 75% of the putative baseline epochs were considered to have a significant change in CS activity (Fig S2a). ***b. Method2***: Consider the CS activity from a 50 ms epoch of one trial as a ‘bin’. We performed a t-test between 50 bootstrapped bins from a given 50 ms epoch and 50 bootstrapped bins from all the epochs across the entire trial period. This gave as a P-value for that epoch, testing the null hypothesis that the activity in that epoch was statistically similar to the activity in any other epoch. We repeated this for every 50 ms epoch and found the epochs that had a P value < 0.05 (Fig S2b). ***c. Method3***: We considered −100 to 0 ms aligned on cue onset as the chosen baseline epoch and compared the activity in that epoch with every other epoch using a conventional approach. The epochs with P<0.05 were considered to have significantly different firing rate from the ‘baseline’ rate (Fig S2c).

**Figure S3:**
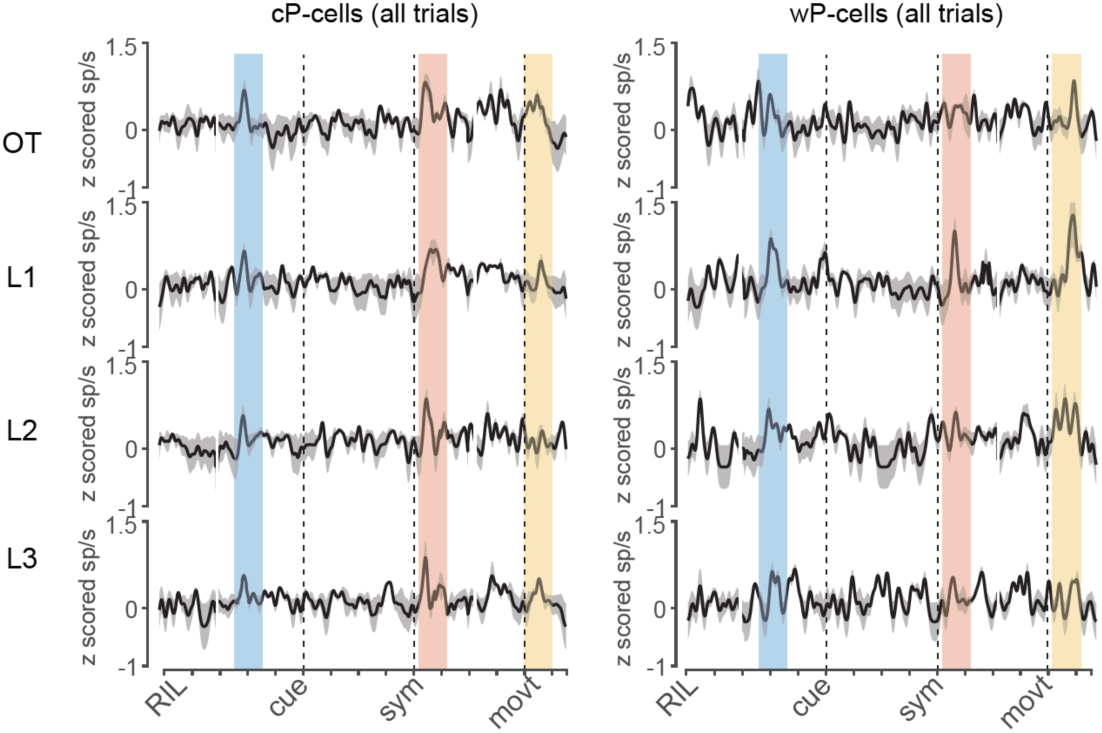
z-scored CS activity for cP-cells and wP-cells. Left: z-scored spike density functions for OT, L,1 L2 and L3 conditions. Right: Same as left but for all wP-cells. Blue, red and yellow shaded regions represent CSx, CSs and CSr epochs respectively.

**Figure S4:**
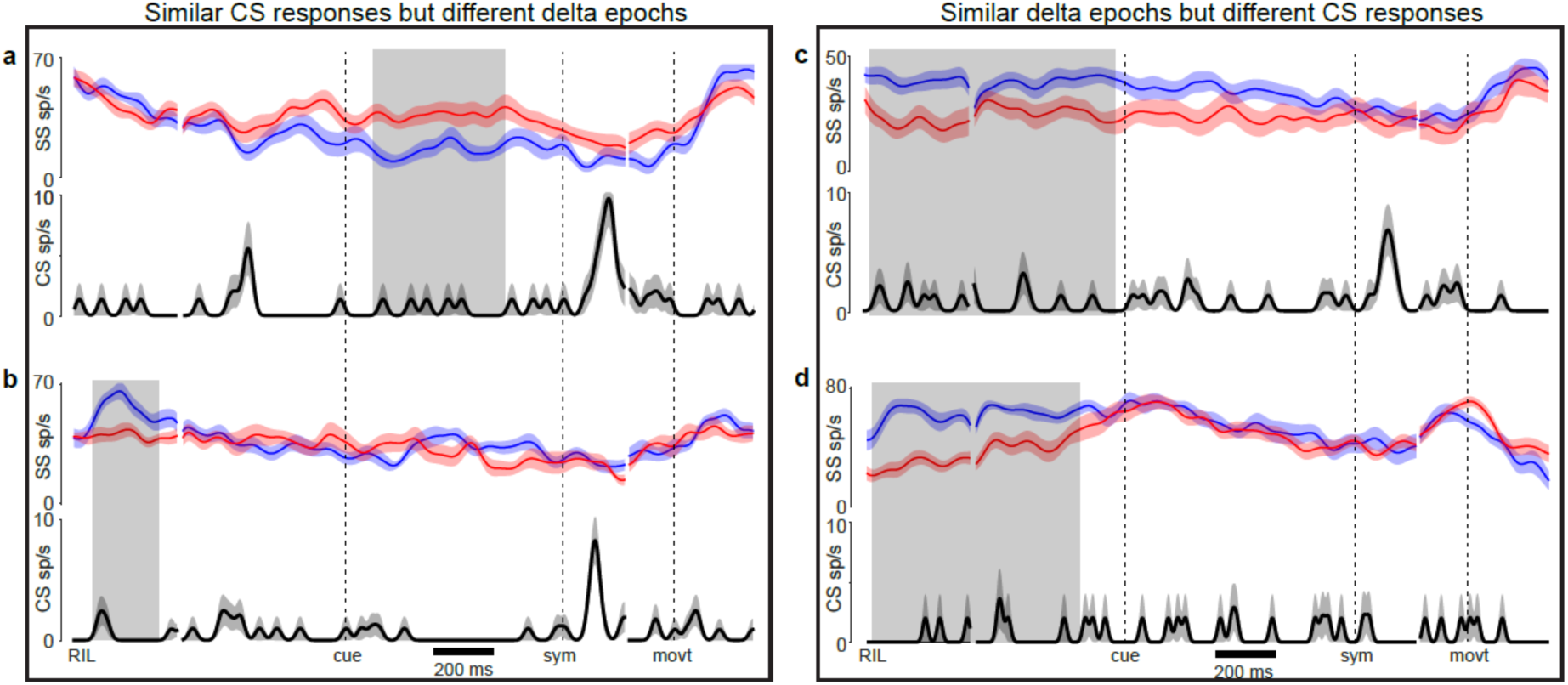
Dissociation between simple spike delta and complex spike responses. a and b: Two P-cells with similar CS responses (increased activity in symbol epoch) but very different SS delta activities. c and d: Two P-cells with similar SS delta activities (similar duration, both cP-cells) but with very different CS responses.

